# Importance of head movements in gaze tracking during table tennis forehand stroke

**DOI:** 10.1101/2023.03.14.532680

**Authors:** Ryosuke Shinkai, Shintaro Ando, Yuki Nonaka, Yusei Yoshimura, Tomohiro Kizuka, Seiji Ono

**Affiliations:** Graduate School of Comprehensive Human Sciences, University of Tsukuba, Ibaraki, Japan; Faculty of Health and Sport Sciences, University of Tsukuba, Ibaraki, Japan

**Author notes:** Corresponding author [ ].

**Keywords:** Gaze, Head movement, VOR, Table tennis, Visual strategy

## Abstract

The purpose of this study was to clarify the properties of gaze and head movements during forehand stroke in table tennis. Collegiate table tennis players (n = 12) conducted forehand strokes toward a ball launched by a skilled experimenter. A total of ten trials were conducted for the experimental task. Horizontal and vertical movements of the ball, gaze, head and eye were analyzed from the image recorded by an eye tracking device. The results showed that participants did not always keep their gaze and head position on the ball throughout the entire ball path. Our results indicate that table tennis players tend to gaze at the ball in the initial ball-tracking phase. Furthermore, there was a significant negative correlation between eye and head position especially in the vertical direction. This result suggests that horizontal VOR is relatively suppressed than vertical VOR in ball-tracking during table tennis forehand stroke. Finally, multiple regression analysis showed that the contribution of head position to gaze position was significantly higher than that of eye position. This result indicates that gaze position during forehand stroke could be associated with head position rather than eye position. Taken together, head movements may play an important role in maintaining the ball in a constant egocentric direction in table tennis forehand stroke.

## 1. Introduction

In table tennis, successful hitting is a greatly difficult task for players as the average number of shots a rally (shots/rally) is only four regardless players participated in the world championships and the Olympic games (Leite et al., 2017). Furthermore, table tennis rallies are characterized by severe time constraints. Therefore, acquiring visual information (mainly a ball) could be one of the most important factors in better understanding spatial and temporal information during table tennis performance.

Although “keep your eyes on the ball” during interceptive performance is a traditional coaching manner, it has been critically discussed in research fields because humans cannot track visual targets moving faster than 70 deg/s (Schalen, 1980). Previous studies have reported that players do not keep their eyes on the entire ball path, in cricket (Land and McLeod, 2000; Croft et al., 2010), baseball (Hubbard and Seng, 1954; Bahill and LaRitz, 1984), table tennis (Ripoll and Fleurance, 1988; Rodrigues et al., 2002; Ishigaki, 2007; Shinkai et al., 2022) and squash (Hayhoe et al., 2012). These results suggest that it is not necessary for players completely keep their gaze on the ball until the impact time. Especially in baseball batting, the time of approximately 150 ms before the impact is not tightly related to hitting accuracy (Higuchi et al., 2016; Gray and Cañal-Bruland, 2018), suggesting that players do not make much use of visual information about the time immediately before the impact for successful hitting. In table tennis, it is necessary to look not only at the ball, but also at the opposite player during games. In particular, information about the racket of the opposite player is important for predicting the ball direction (Piras et al., 2016; Piras et al., 2019). Therefore, table tennis players may quit tracking the approaching ball earlier, even in the task where they only track the ball compared to other intercept sports. Shinkai and colleagues (2022) have reported that table tennis players tended to shift their gaze from the approaching ball earlier during the rally task. However, it is uncertain about the visual strategies of table tennis players using methods without the use of rally tasks. Thus, it is necessary to investigate gaze position relative to ball position for further understanding of the property of gaze tracks in table tennis.

Furthermore, head movements could play important roles in interceptive performance because gaze position is determined by eye-head interaction (Roy and Cullen, 2003; Pallus and Freedman, 2016). As mentioned above, there is a physiological limitation to smoothly pursuing a moving target, but gaze tracking can be improved by combining head and eye movements (Bahill and LaRitz, 1984). Higuchi and colleagues (2018) have reported that baseball players are able to track the ball by using not only eye movements but also head movements, suggesting one of the important roles of head tracking in interceptive performance on fast pitches. As one of the other representative roles of head tracking, Mann and colleagues (2013) suggest that it is important for players to keep their head orientation relative to the egocentricity of the ball in visuomotor tasks. This role would contribute to the acquisition of visual information with respect to a moving ball. However, there is no study that examined these roles of head movements during forehand stroke in table tennis. Therefore, we attempted to examine head position relative to the ball during the ball-tracking phase to determine whether these suggestions apply to head tracking in table tennis.

The vestibulo-ocular reflex (VOR) is a reflective eye movement induced by head movements. In this case, eye movements are induced in the opposite direction to the head movements, which play an important role to keep our gaze on a stational visual target. However, VOR is thought to interrupt keeping gaze position on a moving target. Bahill and RaLitz (1984) have reported that professional baseball batters were good at suppressing horizontal VOR to keep their gaze position on the moving ball. Other studies have revealed that although horizontal VOR occurs in the first half of the ball-tracking, gaze position is tightly aligned to the ball (Kishita et al., 2020a and 2020b). Thus, since there is no consistent evidence regarding the effect of VOR on ball tracking in previous studies, this study focused on VOR and visual strategies in table tennis.

Therefore, the purpose of this study was to clarify the properties of gaze and head movements during forehand stroke in table tennis. In this study, we attempted to quantify not only the horizontal component of gaze and head position, but also their vertical component. We focused on the following three points (1) whether their gaze is kept on the ball throughout the entire ball path, (2) whether the head motion direction is consistent with the ball trajectory, (3) how the horizontal and vertical VOR affects gaze movements in response to head motion.

## 2. Materials and Methods

### 2.1. Participants

The participants were twelve male college students belonging to a table tennis team (mean age: 20.7 ± 1.2 years, height: 168.8 ± 4.0 cm, body mass: 63.2 ± 4.8 kg, table tennis experience: 11.1 ± 2.7 years) and they reported having normal or corrected to normal vision and no known motor deficits. The skill levels of the participants were representatives of the prefecture (8 players) and city (4 players). The representative players of the prefecture had a lot of experience by participating in the annual all Japan table tennis championship. They were diagnosed neither as a stereoscopic problem nor strabismus. All participants gave their informed consent to participate in the experiment. This study was conducted in accordance with the Declaration of Helsinki, and all experimental protocols were approved by Research Ethics Committee at the Faculty of Health and Sport Sciences, University of Tsukuba. Written informed consent was obtained from all participants before their participation.

### 2.2. Experimental procedure

Participants wore an eye-tracking device (Tobii Pro glasses 2, Tobii Technology, Stockholm) to record gaze, ball and head angles during experimental tasks, and the calibration was performed before the experiment. The participant (Fig 1A-➀) conducted the forehand strokes as an experimental task. The experimenter delivered a ball to one target (diameter: 24 cm, Fig 1A-➂) on the participant’s side, and the participant aimed to hit the ball to the circular target (diameter: 32 cm, Fig 1A-➃) on the experimenter’s side. Participants conducted three practice trials to become familiar with this task, followed by a total of ten trials for the experimental task. Please note that they did not keep performing until they had ten successful trials.

**Fig.1.**
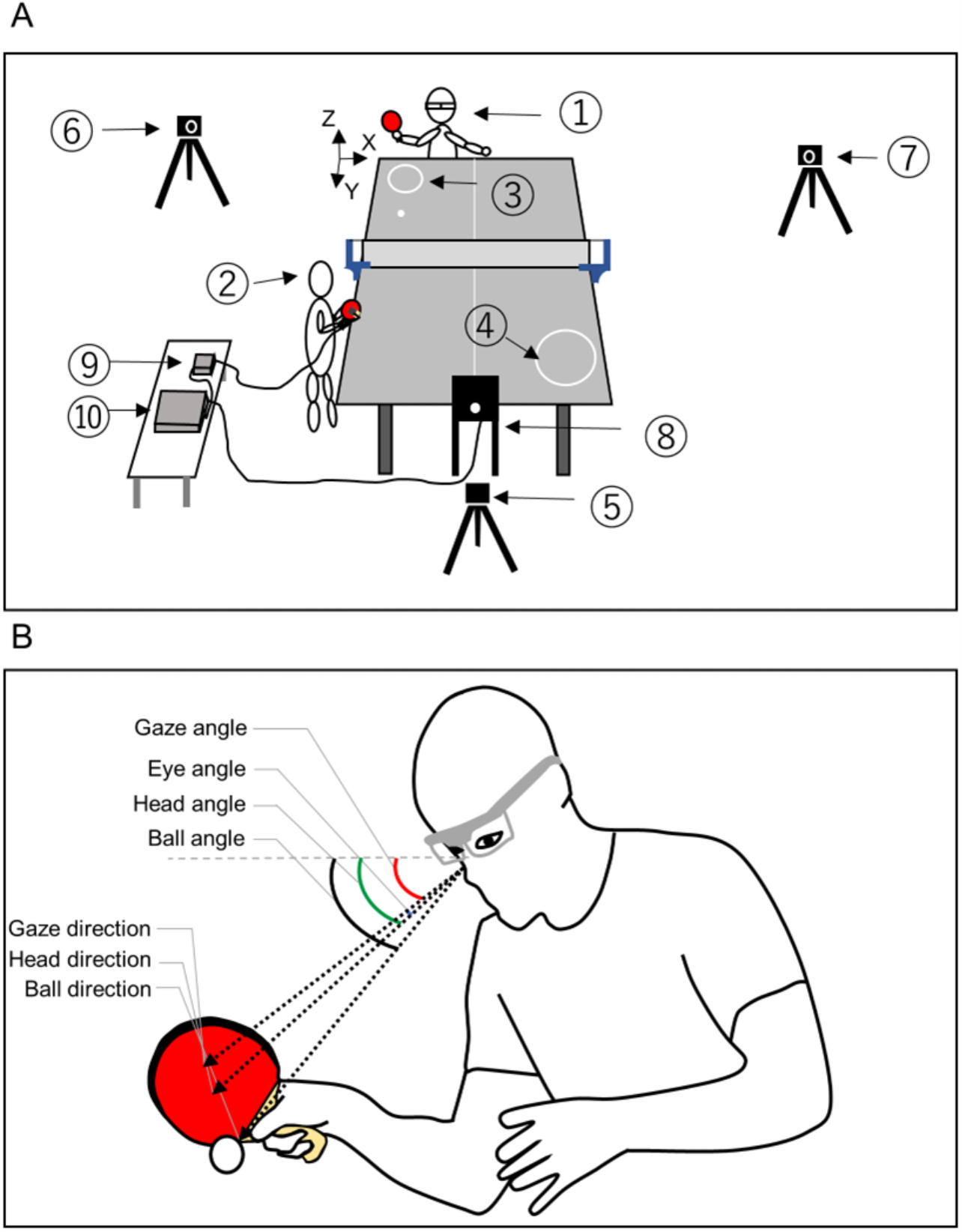
Overview of the experimental task (A) and measurement of the gaze, ball eye and head angles (B). (A) ➀participant, ➁skilled experimenter, ➂participant-side circular target, ➃ experimenter-side circular target, ➄➅➆high speed camera, ➇LED light, ➈control box for gyro sensor, ➉waveform generator. (B) Four angles were calculated based on the video footage of the scene camera of the eye tracking device in each frame.

### 2.3. Data synchronization

To synchronize the impact time of the scene camera of the eye-tracking device and the high-speed camera (Fig 1A-➄, ➅, ➆), an acceleration sensor was attached to the rear of the racket of the experimenter, and the LED lights (Fig 1A-➇) was set to provide the signal output from the acceleration sensor (MP-A0-01A, MicroStone, Japan, Fig 1A-➈) through the waveform generator (SG-4211, IWATU, Japan, Fig 1A-➉).Therefore, the LED lights were flashed by the vibration at the moment of the experimenter’s impact. The delay from the hitting time to LED flash was <10 ms. The time when the front LED light flashed was captured from each image of the high-speed camera (frame rate =240 fps, EX-ZR200, CASIO, Japan, Fig 1A-➅, ➆). The high-speed camera (Fig 1A-➄) was set to reflect the rear LED flash (Fig 1A-➉) meant the stroke movement of participants to determine the impact time. To synchronize these data, the time of each trial was normalized so that Time=0 was at the stroke of the experimenter and 100% was at the return of the participant. The images from the scene camera of the eye-tracking device were recorded at a sampling frequency of 25 Hz.

### 2.4. Data analysis

Video footage from the scene camera of the eye-tracking device was digitized using motion analysis software (Frame-DIAS IV, DKH, Japan) to determine the coordinates (pixels) of three spatial positions of the ball, eye and top-right corner of the table tennis net in each frame of the video footage. The coordinates of the top-right table tennis net position were used to calculate for head rotation to ensure that all three angles were reported relative to global coordination. The coordinates on the footage of the scene camera were digitized as x and y values. The changes of digitized the x values from the top-right net position reflect the head rotation of the yaw while the y values reflect the pitch. The zero point was each data (gaze/ eye/ head/ ball) at the moment of ball launch (the first digitized point in each trial). The coordinates of these positions were used to calculate the ball, gaze, head and eye angle (deg) subtended at each data relative to the initial direction at the moment of ball-launch (Fig 1B). The pixel values of the coordinates were converted to angles based on the specifications of the eye-tracking device (horizontal angle: 82 deg / 1920 px, vertical angle: 52 deg / 1080 px). Three relative angles were calculated to convey the comparative position of the three raw angles: the gaze, head and eye angles relative to the ball angle.

We counted the number of successful shots that participants hit the aimed target to assess the performance of the shots. We used a high-speed camera (Fig 1A-➆) to count this number.

### 2.5. Statistical analysis

To examine the differences in the angle between gaze and ball, between head and ball at each normalized time, an independent t-test was performed. To examine a pattern of ball, gaze, eye and head shifts, a one-way ANOVA was performed to analyze gaze, eye and head positions in each direction. To examine the relationship between eye and head position corresponding to each same time, the Pearson correlation coefficient was calculated in each direction.

To examine the contribution of eye and head position to gaze position, a multiple regression analysis using stepwise regression was conducted with gaze position as the dependent variable and eye and head position as the independent variables. When horizontal gaze position was the dependent variable, horizontal eye and head position were the independent variables, and when vertical gaze position was the dependent variable, vertical eye and head position were the independent variables.

An alpha level of 5 % was applied for all the tests. All statistical tests were conducted by IBM SPSS software version 27 (SPSS Inc, USA). Unless noted otherwise, data are presented as mean ± standard deviation (SD).

### 2.6. Properties of ball trajectories hit by the experimenter

We analyzed ball trajectories hit by the experimenter to confirm the reliability of the test. For the ball trajectory, we calculated the position and velocity in all trials by using motion analysis software (Frame-DIAS IV, DKH, Japan). Two high-speed cameras (frame rate =240 fps, EX-ZR200, CASIO, Japan, Fig 1A-➅, ➆) were used for capturing ball trajectories. The origin of the coordinate system was the position of the floor directly below the corner of the table tennis table (Fig 1A).

Figure 2 shows the ball trajectories in three-dimensional geometry. All trajectories are consistent throughout the experiment. Furthermore, figure 3 shows the mean ball position and velocity trajectories in each coordinate. Supplemental Figure 1 shows all data (120 traces) used to calculate the mean ± SD in each coordinate. The time throughout the ball trajectories was 505.5 ± 27.1 ms.

**Fig.2.**
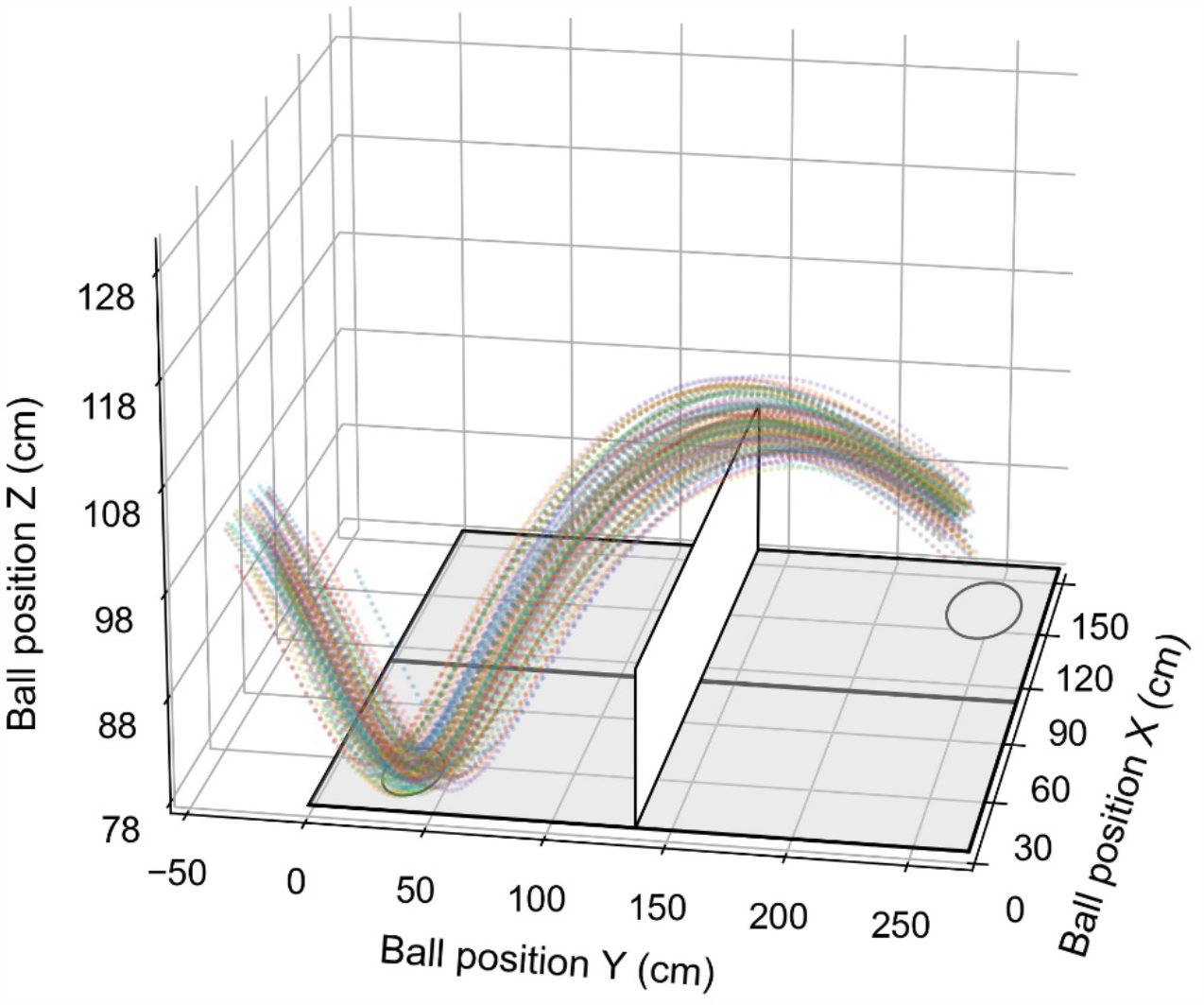
Three-dimensional ball trajectories of the shots by the experimenter in all trials. Each trace shows one ball trajectory during a trial. The grey rectangle parallel to the XZ-plane shows the net of the table tennis table. The circle on the XY-plane shows the circle that the experimenter aimed to hit the ball as the first ball-bounce area.

**Fig.3.**
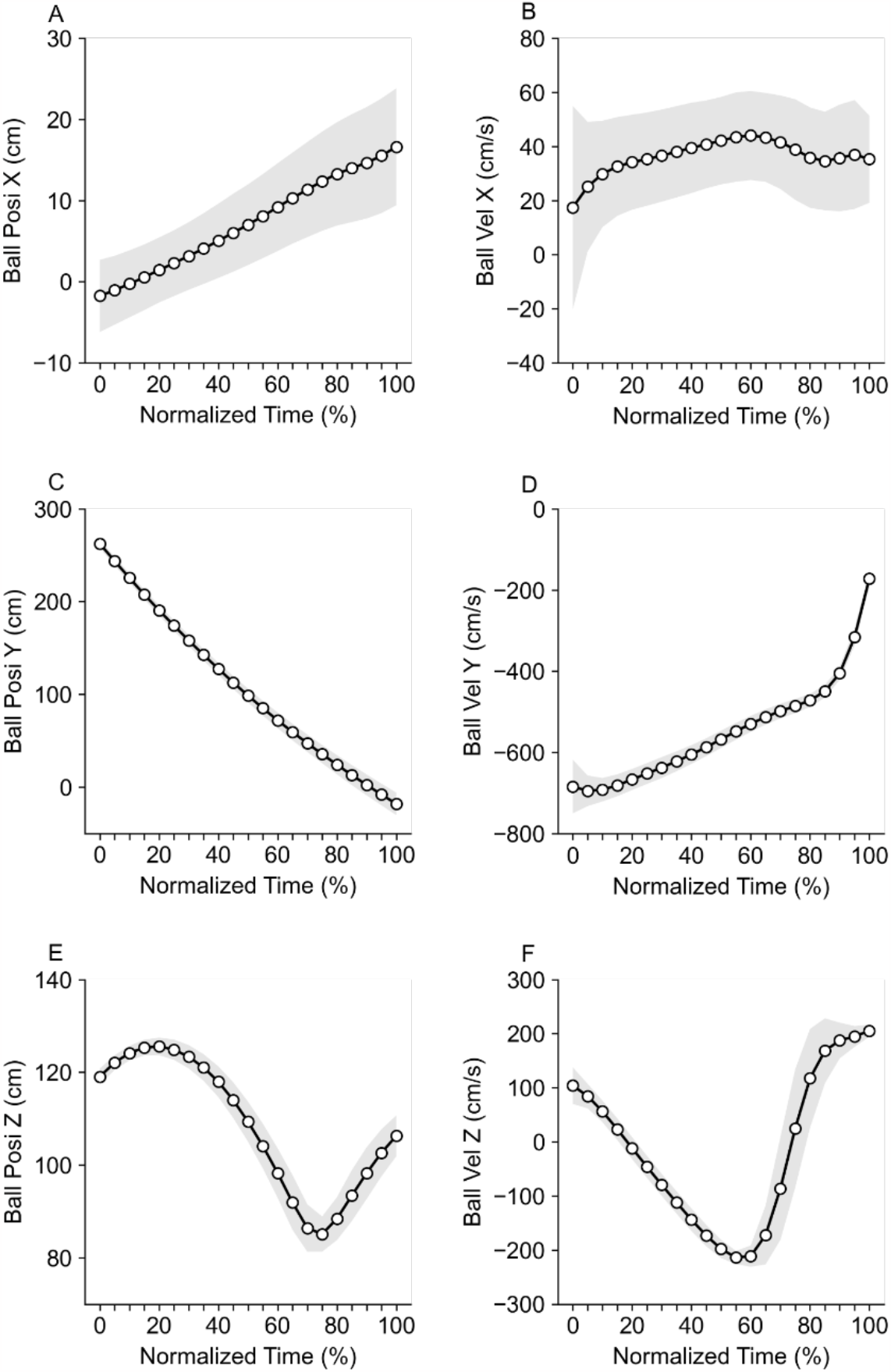
Mean ball positions and velocities of the shots by the experimenter as a function of normalized time in X (A, B), Y (C, D) and Z (E, F) coordinates. Grey shaded areas show standard deviation.

To examine the effect of ball trajectories on the main results, the Pearson correlation coefficient was calculated to examine the correlation between ball trajectory (position and velocity) and gaze, head positions in X- and Z-coordinates. Supplemental Table 1 shows the results at each normalized time, indicating that there are no significant correlations between ball trajectories and main results.

## 3. Results

### 3.1. Gaze and ball position in the ball-tracking phase

Independent t-test for gaze and ball position in the horizontal direction showed that horizontal gaze position was significantly smaller (more leftward) than horizontal ball position at 0-10, 10-20, 70-80, 80-90 and 90-100 % of the time. Furthermore, there were significant differences between gaze and ball shifts (gaze: 0-10 vs 10-20, 10-20 vs 20-30, 20-30 vs 30-40, 30-40 vs 40-50, 40-50 vs 50-60, 50-60 vs 60-70, 60-70 vs 70-80 %, ball: 0-10 vs 10-20, 10-20 vs 20-30, 20-30 vs 30-40, 30-40 vs 40-50, 40-50 vs 50-60, 50-60 vs 60-70, 60-70 vs 70-80, 70-80 vs 80-90, 80-90 vs 90-100 %, *p* < 0.01, Fig 4A).

**Fig 4.**
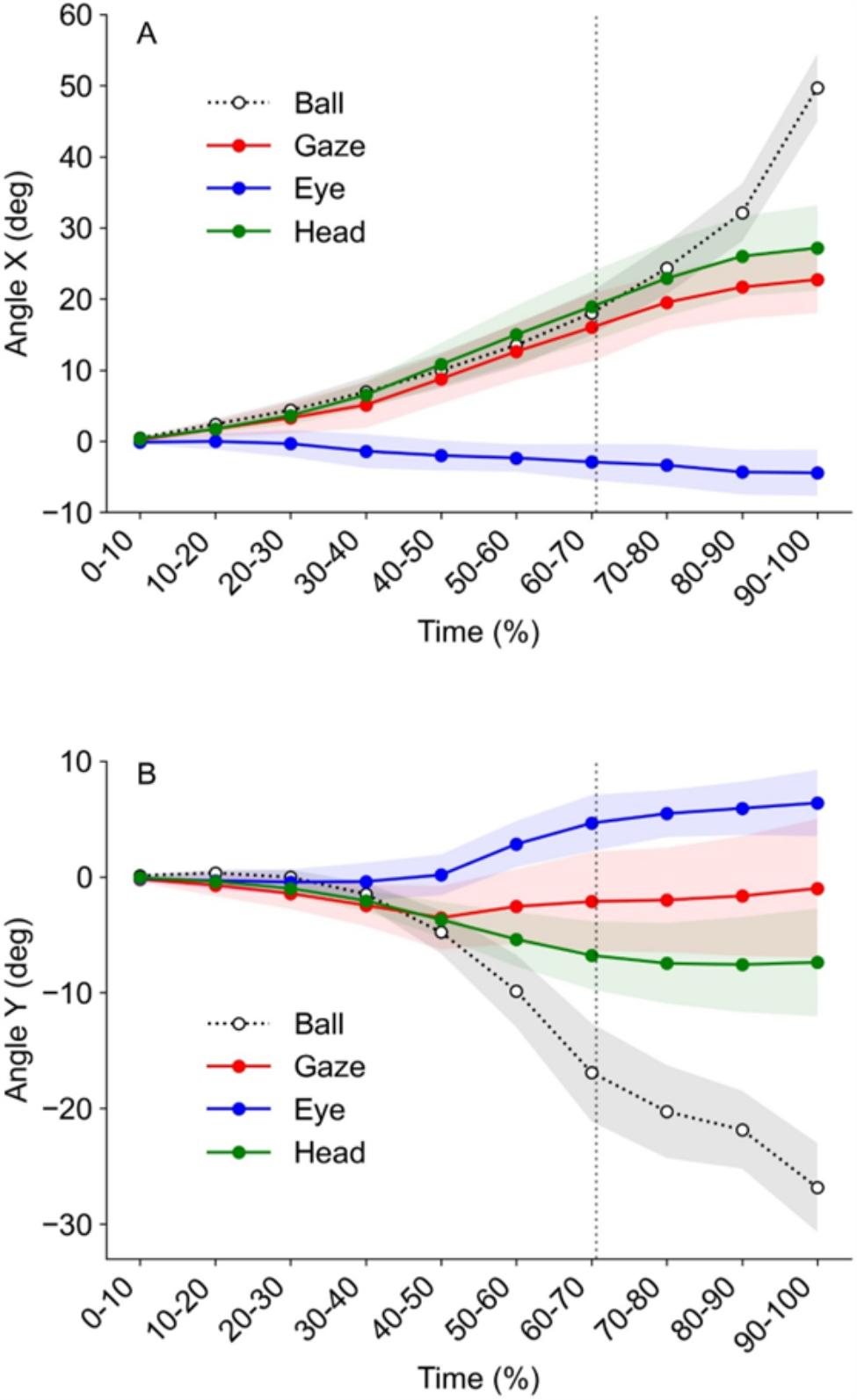
Ball (white), gaze (red), eye (blue) and head (green) position in the horizontal (A) and vertical (B) directions. Dotted vertical lines show the bounce time of the launched ball by the experimenter. Shaded areas represent standard deviation in each parameter.

Independent t-test for gaze and ball position in the vertical direction showed that vertical gaze position was significantly smaller (more upward) than vertical ball position at 0-10, 10-20, 20-30, 50-60, 60-70, 70-80, 80-90 and 90-100 % of the time. Furthermore, there were significant differences between gaze and ball shifts (gaze: 0-10 vs 10-20, 10-20 vs 20-30, 20-30 vs 30-40, 30-40 vs 40-50, 40-50 vs 50-60, 50-60 vs 60-70, 60-70 vs 70-80 %, ball: 0-10 vs 10-20, 10-20 vs 20-30, 20-30 vs 30-40, 30-40 vs 40-50, 40-50 vs 50-60, 50-60 vs 60-70, 60-70 vs 70-80, 70-80 vs 80-90, 80-90 vs 90-100 %, *p* < 0.01, Fig 4B).

Figure 5 shows the gaze position relative to the ball position in each normalized time point. The gaze positions relatively shifted left- and upward relative to the ball position as time proceeded. These plots are consistent with the results described above.

**Fig 5.**
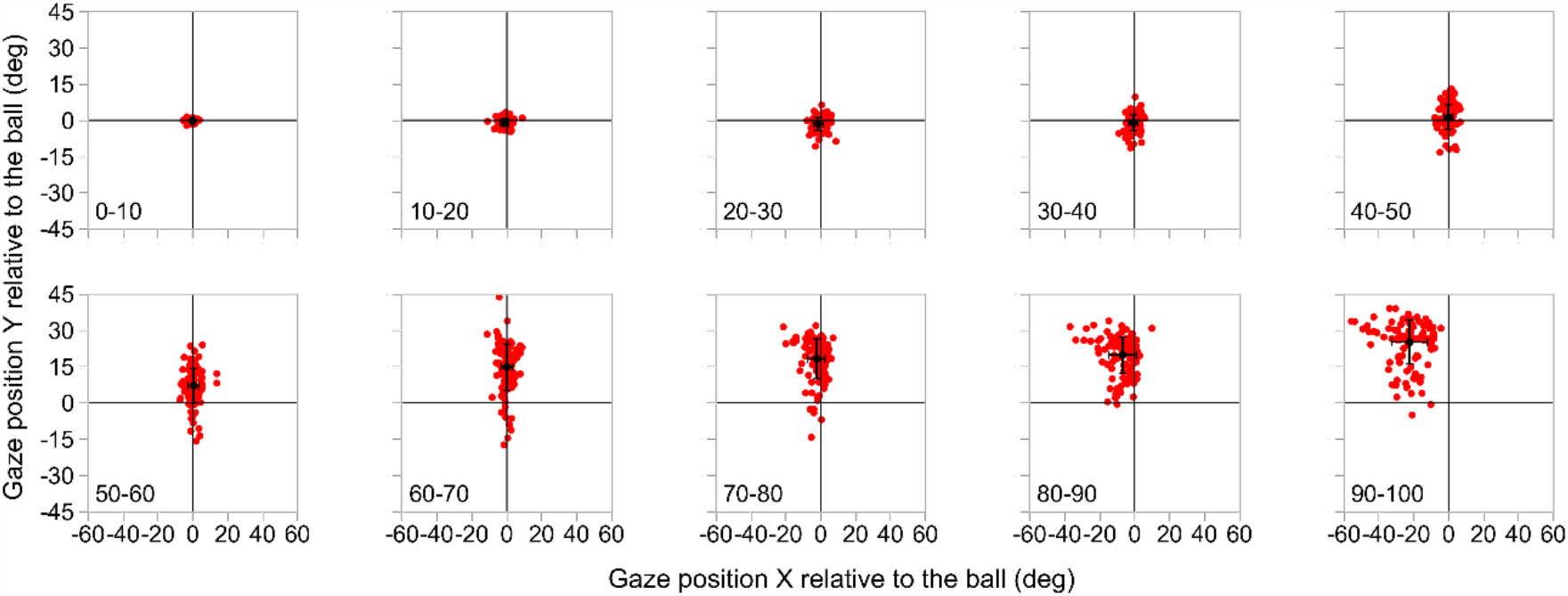
Gaze position relative to the ball position in each normalized time point. Each red dot shows one trial data at each time (120 trials in total). Black dots attached to horizontal and vertical bars show the mean ± SD of all plots at each time point.

### 3.2. Head and ball position in the ball-tracking phase

Independent t-test for head and ball position in the horizontal direction showed that horizontal head position was significantly smaller (more leftward) than horizontal ball position at 0-10, 10-20, 80-90 and 90-100 % of the time. Furthermore, there were significant differences between head and ball shifts (head: 0-10 vs 10-20, 10-20 vs 20-30, 20-30 vs 30-40, 30-40 vs 40-50, 40-50 vs 50-60, 50-60 vs 60-70, 60-70 vs 70-80 %, ball: 0-10 vs 10-20, 10-20 vs 20-30, 20-30 vs 30-40, 30-40 vs 40-50, 40-50 vs 50-60, 50-60 vs 60-70, 60-70 vs 70-80, 70-80 vs 80-90, 80-90 vs 90-100 %, *p* < 0.01, Fig 4A).

Independent t-test for head and ball position in the vertical direction showed that vertical head position was significantly smaller (more upward) than vertical ball position at 0-10, 10-20, 20-30, 50-60, 60-70, 70-80, 80-90 and 90-100 % of the time. Furthermore, there were significant differences between head and ball shifts (head: 0-10 vs 10-20, 10-20 vs 20-30, 20-30 vs 30-40, 30-40 vs 40-50, 40-50 vs 50-60, 50-60 vs 60-70, 60-70 vs 70-80 %, ball: 0-10 vs 10-20, 10-20 vs 20-30, 20-30 vs 30-40, 30-40 vs 40-50, 40-50 vs 50-60, 50-60 vs 60-70, 60-70 vs 70-80, 70-80 vs 80-90, 80-90 vs 90-100 %, *p* < 0.01, Fig 4B).

Figure 6 shows the head position relative to the ball position in each normalized time point. The head positions shifted left- and upward relative to the ball position as time proceeded. These plots are consistent with the results described above.

**Fig 6.**
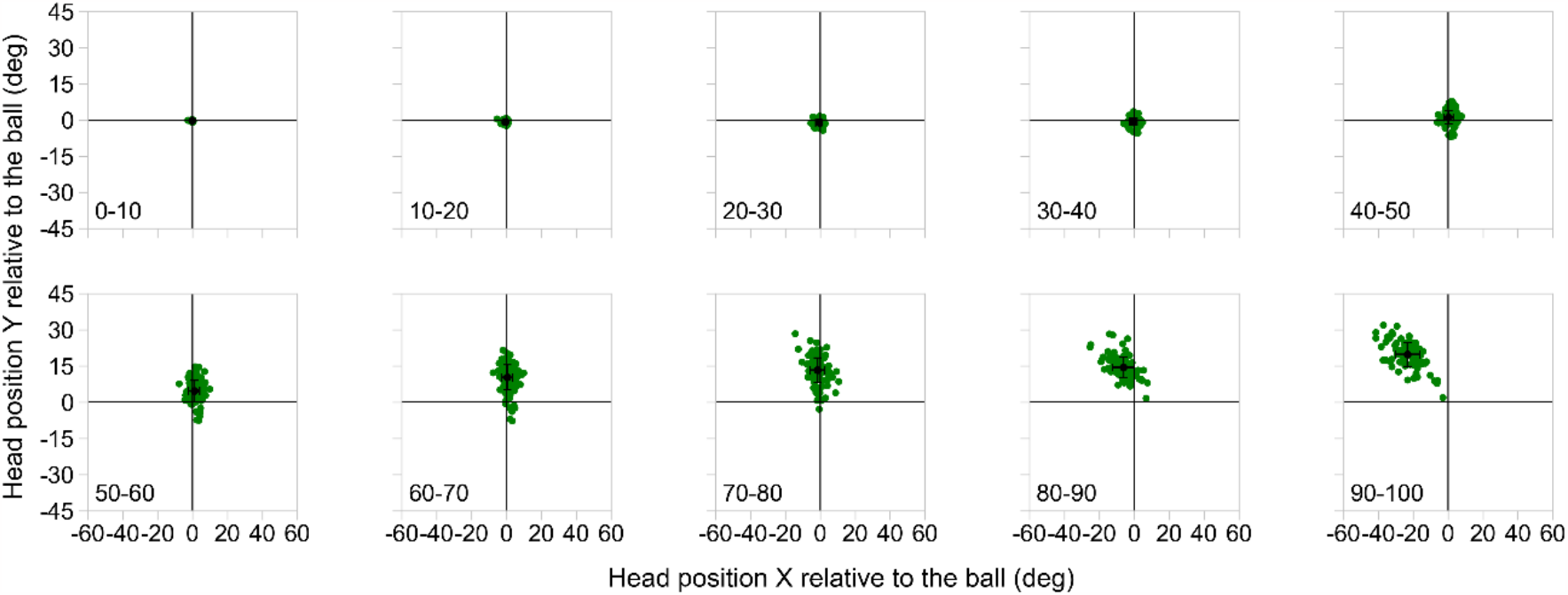
Head position relative to the ball position in each normalized time point. Each green dot shows one trial data at each time (120 trials in total). Black dots attached to horizontal and vertical bars at each time show the mean ± SD of all plots at each time point.

### 3.3. Eye positions relative to the head position in the ball-tracking phase

There was a significant main effect of time on eye position in horizontal direction (*F*_(9, 99)_ = 11.1, *p* < 0.01, *η*^2^ = 0.50, power = 1.0), and significant differences in the post-hoc test (0-10 vs 80-90 and 90-100 %, 10-20 vs 50-60, 60-70, 70-80, 80-90 and 90-100 %, 20-30 vs 50-60, 60-70, 80-90 and 90-100%, *p* < 0.05, Fig 4A). The results indicate that eye position in the second half of the time was significantly smaller (more leftward) than that in the first half of the time.

There was a significant main effect of time on the eye position in the vertical direction (*F*_(9, 99)_ = 44.1, *p* < 0.01, *η*^2^ = 0.80, power = 1.0), and significant differences in the post-hoc test (0-10 vs 50-60, 60-70, 70-80, 80-90 and 90-100 %, 10-20 vs 50-60, 60-70, 70-80, 80-90 and 90-100 %, 20-30 vs 50-60, 60-70, 70-80, 80-90 and 90-100%, 30-40 vs 50-60, 60-70, 70-80, 80-90 and 90-100%, 40-50 vs 60-70, 70-80, 80-90, 90-100%, 50-60 vs 60-70, 70-80, 80-90, 90-100%, *p* < 0.05, Fig 4B). The results indicate that eye position in the second half of the time was significantly larger (more upward) than that in the first half of the time.

These results indicate that the change in eye position was in the opposite direction of the head position in each direction (Fig 4). Figure 7 shows eye position relative to head position, indicating that the eye position shifted in the opposite direction to head position in the last phase, mainly in the vertical direction. For example, downward head motion produces an upward VOR, as shown in figure 4B. Therefore, this result suggests that typical VOR occurs in the vertical direction, whereas VOR could be suppressed in the horizontal direction.

**Fig 7.**
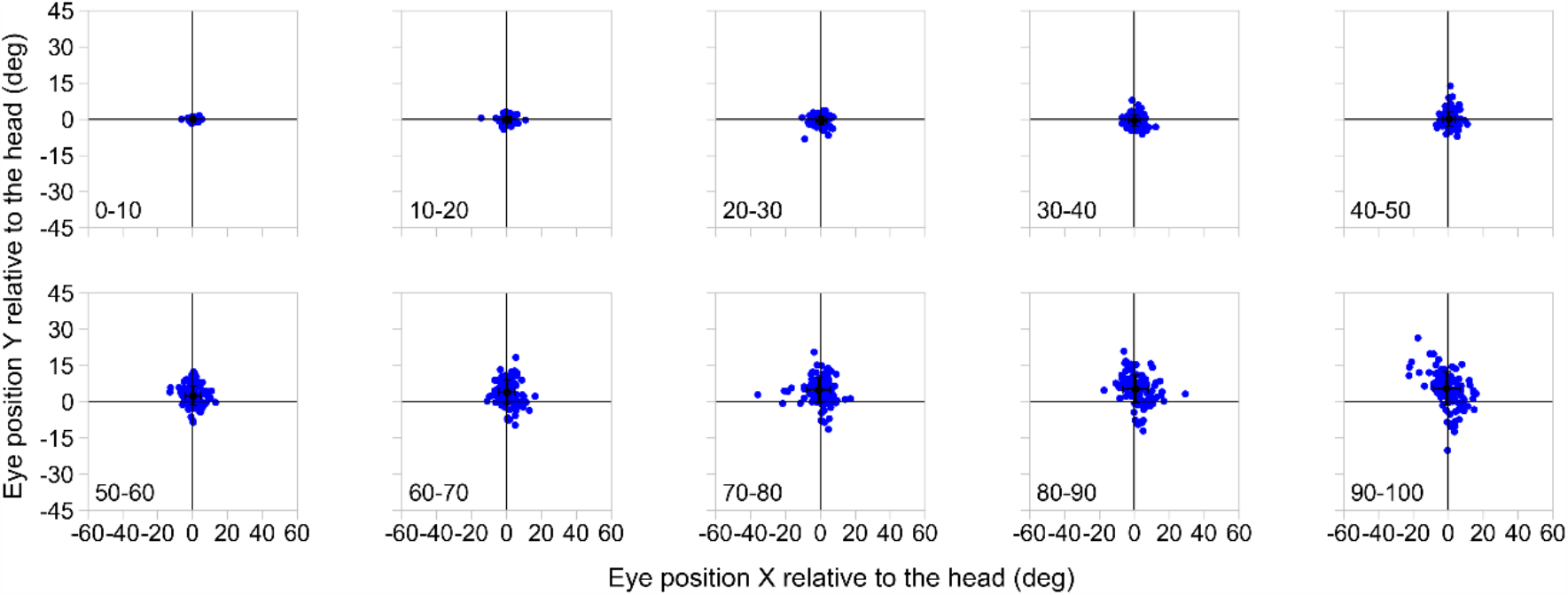
Eye position relative to the head position in each normalized time point. Each blue dot shows one trial data at each time (120 trials in total). Black dots attached to horizontal and vertical bars at each time show the mean ± SD of all plots at each time point.

### 3.4. Relationship between eye and head position in the ball-tracking phase

The Pearson correlation between eye and head position was examined with 120 data (10 data a participant) in the horizontal (A) and vertical (B) directions (Fig. 8). A significant correlation was found in each direction (horizontal: *r* = -0.53, *p* < 0.01, Fig 8A, vertical: *r* = -0.76, *p* < 0.01, Fig 8B).

**Fig 8.**
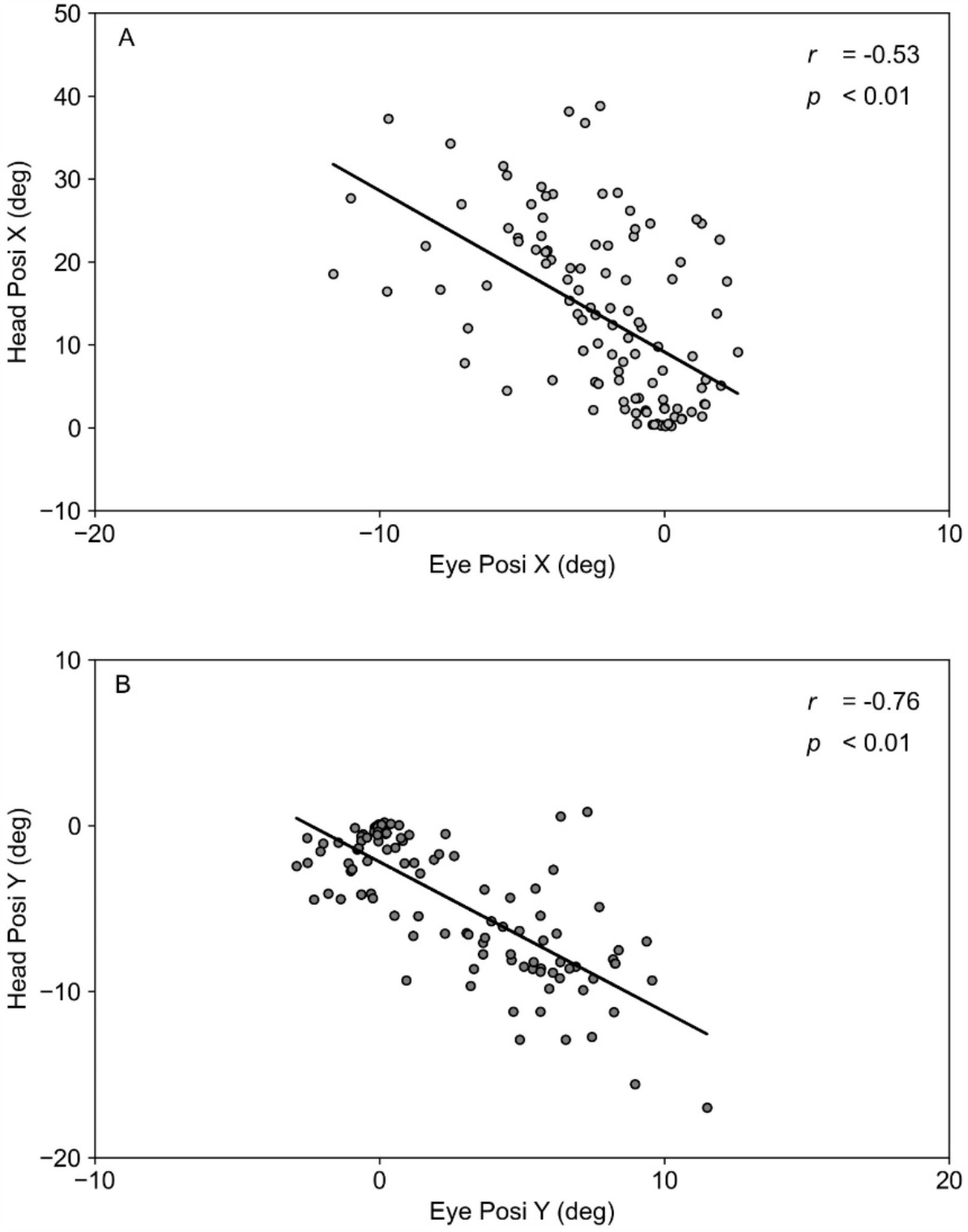
Relationship between eye and head position. The Pearson correlation between eye and head position was examined with 1,200 data (10 data a trial) in the horizontal (A) and vertical (B) directions. A significant negative correlation was found in each direction (horizontal: *r* = -0.23, *p* < 0.01, Fig 8A, vertical: *r* = -0.64, *p* < 0.01, Fig 8B).

### 3.5. Contribution of eye and head position relative to gaze position in the ball-tracking phase

The multiple regression analysis described significant regression equations. There were significant positive correlations between gaze and eye position, and between gaze and head position (Fig 9A-9D). The valuables of eye position in both directions were excluded from equations by stepwise regression. The standardized coefficients (β) are shown in Table 1. The standardized coefficients of head position were significantly higher than that of eye position. The VIFs were all <10.0 and there was no problem with multicollinearity. These results indicate that the effect of head position on the gaze position is significantly higher than that of eye position.

**Fig 9.**
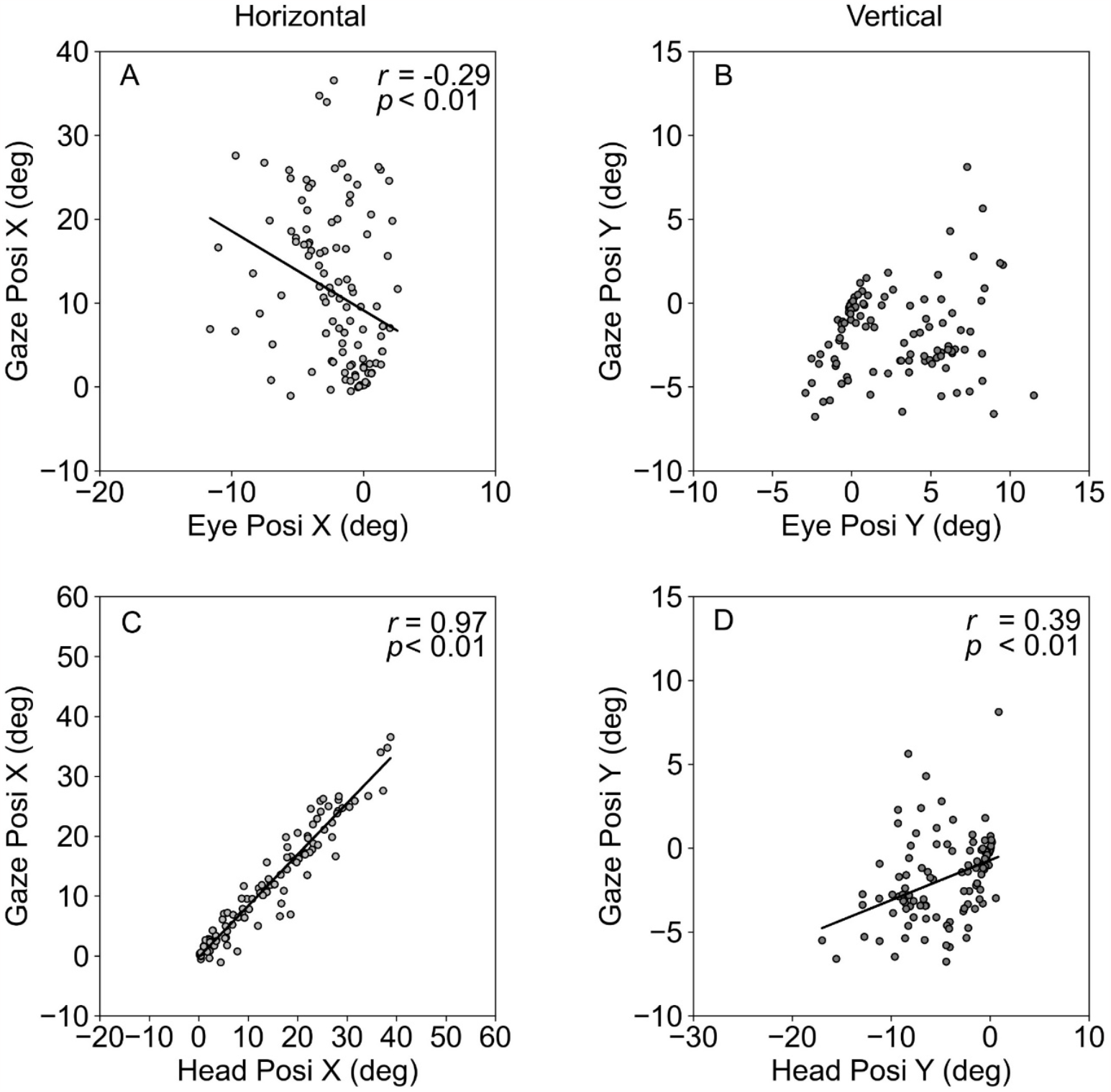
Relationship between gaze and eye position, between gaze and head position in the horizontal and vertical direction. The left two panels show the horizontal data between gaze and eye position (A), and between gaze and head position (C). The right two panels show the vertical data between gaze and eye position (B), and between gaze and head position (D).

**Table 1.**

| A |  |  |  |  |  |  |
| --- | --- | --- | --- | --- | --- | --- |
|  | Dependent Valueable | Independent Valueable | Adjusted R<sup>2</sup> | DOF | *F* | *p* |
| Horizontal | Gaze Posi | Eye Posi<br>Head Posi | 0.86 | 1, 198 | 7411.6 | < 0.01 |
| Vertical | Gaze Posi | Eye Posi<br>Head Posi | 0.52 | 1, 198 | 1308.0 | < 0.01 |
| B |  |  |  |  |  |  |
| | Dependent Valueable | Independent Valueable | $\beta$ | *t* | *p* | VIF |
| Horizontal | Gaze Posi | Head Posi | 0.93 | 86.1 | < 0.01 | 1.0 |
| Vertical | Gaze Posi | Head Posi | 0.72 | 36.2 | < 0.01 | 1.0 |

### 3.6. Hitting accuracy

The number of successful hitting was 6.8 ± 1.0 shots. We separated the gaze position into two groups (Hit and Miss), and independent t-test showed that there was no significant difference at each normalized time (Fig 10).

**Fig 10.**
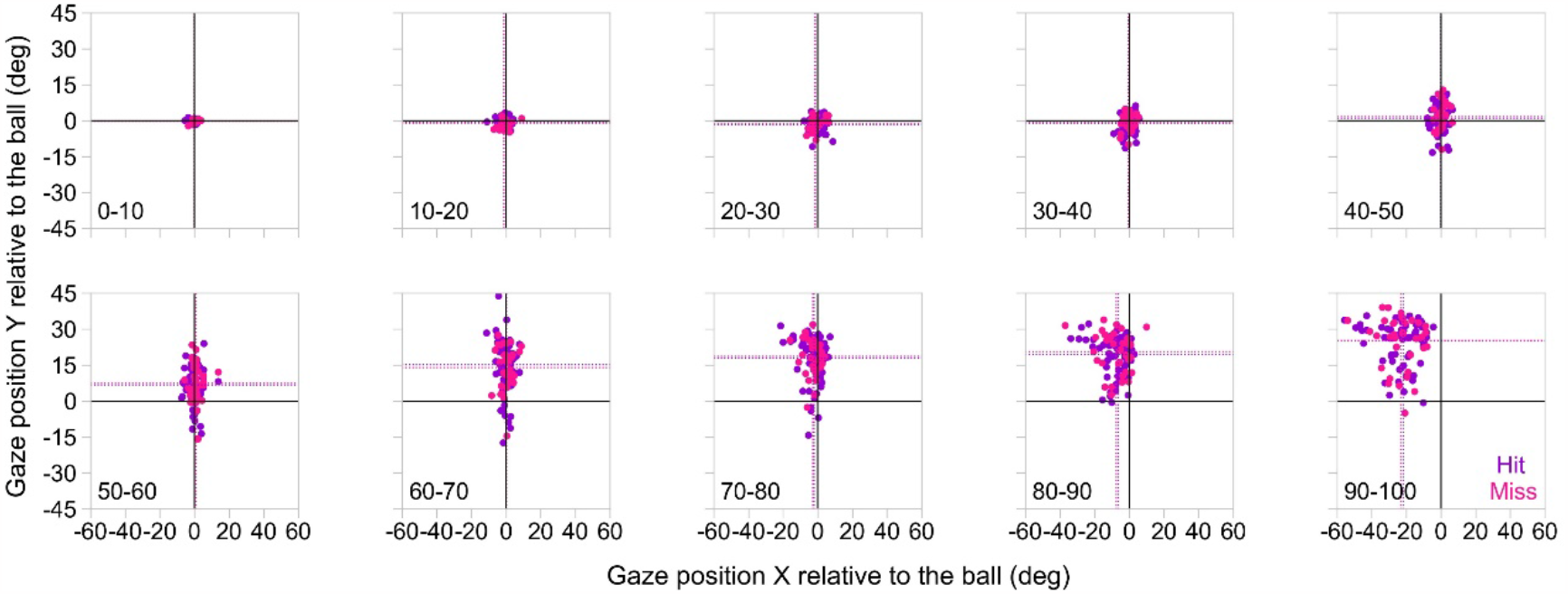
Gaze position relative to the ball position in each normalized time point. Violet dots indicate single trial data for hit at each time point, whereas pink dots indicate single trial data for misses at each time point. The horizontal and vertical dotted lines indicate the mean values of at each time point.

## 4. Discussion

In the present study, we examined the properties of gaze and head movements for table tennis players while they conducted a forehand stroke task. We found that participants did not always keep their gaze and head position on the ball position throughout the entire ball path. Our result indicate that table tennis players tend to gaze at the ball in the initial ball-tracking phase. Here, we discuss the properties of gaze, head and eye movements relative to the ball during the table tennis forehand stroke for further understanding of visual strategies in table tennis.

### 4.1. Gaze position relative to the ball position

The horizontal gaze position was lagging behind the ball position at 70-80, 80-90 and 90-100 % of the time. The vertical gaze position was also lagging behind the ball position at 50-60, 60-70, 70-80, 80-90 and 90-100 % of the time. These results indicate that the participants in the present study did not always keep their gaze positions on the ball throughout the entire ball trajectory, which partly supports the report by Ripoll and Fleurance (1988) showing that table tennis players tracked a certain part of the ball path rather than the entire flight path. Our results are also consistent with previous studies by Rodrigues et al (2002), Ishigaki (2007) and Shinkai et al (2022). They have clarified the visual strategies during table tennis rallies, showing that table tennis players shift their eyes from the ball in the earlier phase of ball-tracking. Therefore, our results suggest that it is important for table tennis players to track the ball in the first half of ball trajectories.

Ripoll and Fleurance (1988) have reported that the gaze is finally directed to the ball at the time of ball-racket contact for only five table tennis players. However, this finding does not generalize when averaged over the population of table tennis players (n = 12) tested in this study. It has also been revealed that even though the gaze is not directed to the ball immediately before the bat-ball contact (approximately 150 ms), intercept performance is not decreased in comparison with when the gaze was kept on the ball throughout the entire ball path (Higuchi et al., 2016; Gray and Cañal-Bruland, 2018). Given the average time of the entire ball trajectory (505.5 ± 27.1 ms) and the percentage of time the ball bounced (70.6 ± 6.6 %) in this study, it is estimated that the time from the ball bounced to racket-ball contact is approximately 150 ms. Therefore, it is most likely that participants did not make much use of visual information of the ball position for successful performance during this time.

### 4.2. Head position relative to the ball position

The horizontal head position changed leftward relative to the ball position at 80-90 and 90-100 % of the time. On the other hand, the vertical head position changed upward relative to the ball position from 50-60 to 90-100 % of the time range. Although these results are similar to that of the gaze position, they indicate that the horizontal head position is more aligned with the ball position than the horizontal gaze position. Mann and colleagues (2013) suggest that tracking an approaching ball with the head is helpful because it maintains the ball in a constant egocentric direction. Therefore, it is possible that participants in the present study made great use of head tracks to maintain the ball in a constant egocentric direction.

### 4.3. Occurrence of VOR during ball-tracking

The vertical head position moved downward while the vertical eye position moved in the opposite upward direction. In addition, there was higher anti-correlation between eye and head position in the vertical direction. These results reflect the occurrence of typical VOR. On the other hand, in the horizontal direction, eye position showed a slight change relative to the rightward change in head position. In addition, there was lower anti-correlation between eye and head position in the horizontal direction. These results suggest that horizontal VOR is relatively suppressed than vertical VOR. Our findings are consistent with a previous study by support Bahill and RaLitz (1984), demonstrating that the expert batters suppress horizontal VOR to maintain the gaze on a moving ball. Therefore, VOR suppression in the horizontal direction could contribute to maintaining the gaze position on the ball. Another study has reported that gaze position relative to the ball is kept when horizontal VOR occurs in the first half during ball-tracking (Kishita et al., 2020a and 2020b). Thus, future studies would be necessary to clarify how VOR contributes to gaze control during target-tracking.

### 4.4. Contribution of eye and head position to gaze position during ball-tracking

The contribution of head position was significantly higher than that of eye position as the result of multiple regression analysis. This result suggests that ball-tracking with the head is necessary in table tennis forehand stroke. Although the importance of head tracking has been reported in cricket (Mann et al., 2013; Sarpeshkar et al., 2017), this is the first study to demonstrate the importance of head movement during forehand stroke in table tennis.

## 5. Conclusion

This study examined the properties of gaze and head position during table tennis forehand stroke. The results showed that participants did not always keep their gaze and head position on the ball throughout the entire ball trajectory. We also found that horizontal VOR was relatively suppressed than vertical VOR during ball-tracking. Furthermore, the contribution of head position on the gaze position was significantly higher than that of eye position. Our findings suggest that gaze position is tightly associated with head position. Taken together, head movements may play an important role in maintaining the ball in a constant egocentric direction in table tennis forehand stroke.

