## supplemental materials for "Importance of head movements in gaze tracking during table tennis forehand stroke"

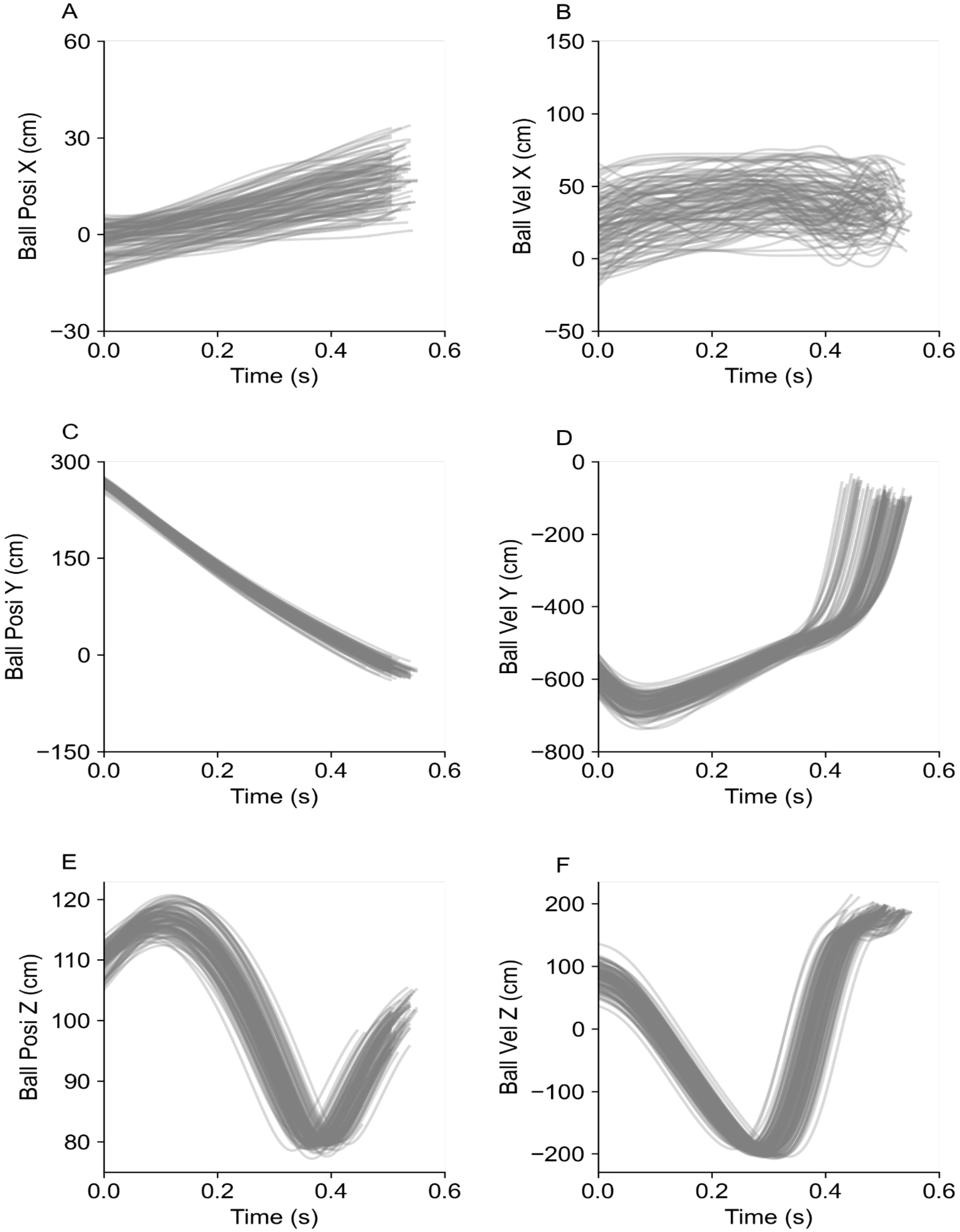


**Supplemental fig 1. All traces of ball positions and velocities in each coordinate.**

**Supplemental Table 1. Pearson correlation between ball trajectories and main results in each normalized time.**

| Normalized Time (%) | | | *r* | *p* |
|---|---|---|---|---|
| 0-10 | Ball Posi X | Gaze X | -0.05 | 0.87 |
| | | Head X | -0.04 | 0.89 |
| | | Eye X | -0.20 | 0.53 |
| | | Gaze-ball X | -0.19 | 0.56 |
| | | Head-ball X | 0.17 | 0.61 |
| | Ball Posi Z | Gaze Y | 0.56 | 0.06 |
| | | Head Y | 0.40 | 0.20 |
| | | Eye Y | -0.09 | 0.78 |
| | | Gaze-ball Y | 0.53 | 0.90 |
| | | Head-ball Y | -0.33 | 0.29 |
| | Ball Vel X | Gaze X | 0.04 | 0.89 |
| | | Head X | -0.01 | 0.98 |
| | | Eye X | 0.08 | 0.80 |
| | | Gaze-ball X | -0.05 | 0.87 |
| | | Head-ball X | -0.26 | 0.42 |
| | Ball Vel Z | Gaze Y | -0.57 | 0.06 |
| | | Head Y | -0.38 | 0.22 |
| | | Eye Y | -0.36 | 0.25 |
| | | Gaze-ball Y | -0.46 | 0.13 |
| | | Head-ball Y | -0.15 | 0.65 |
| 10-20 | Ball Posi X | Gaze X | 0.00 | 0.99 |
| | | Head X | 0.07 | 0.83 |
| | | Eye X | -0.09 | 0.77 |
| | | Gaze-ball X | -0.10 | 0.77 |
| | | Head-ball X | 0.05 | 0.88 |
| | Ball Posi Z | Gaze Y | 0.44 | 0.12 |
| | | Head Y | 0.36 | 0.25 |
| | | Eye Y | -0.37 | 0.23 |
| | | Gaze-ball Y | 0.45 | 0.16 |
| | | Head-ball Y | 0.00 | 0.99 |
| | Ball Vel X | Gaze X | 0.05 | 0.87 |
| | | Head X | 0.05 | 0.89 |
| | | Eye X | -0 | 0.99 |
| | | Gaze-ball X | -0.24 | 0.46 |
| | | Head-ball X | 0.04 | 0.91 |
| | Ball Vel Z | Gaze Y | -0.55 | 0.06 |
| | | Head Y | -0.28 | 0.39 |
| | | Eye Y | 0.36 | 0.25 |
| | | Gaze-ball Y | -0.50 | 0.10 |
| | | Head-ball Y | -0.22 | 0.50 |
| 20-30 | Ball Posi X | Gaze X | -0.02 | 0.94 |
| | | Head X | -0.03 | 0.92 |
| | | Eye X | -0.28 | 0.38 |
| | | Gaze-ball X | -0.20 | 0.53 |
| | | Head-ball X | 0.05 | 0.88 |
| | Ball Posi Z | Gaze Y | 0.37 | 0.24 |
| | | Head Y | 0.34 | 0.29 |
| | | Eye Y | 0.37 | 0.24 |
| | | Gaze-ball Y | 0.44 | 0.16 |
| | | Head-ball Y | -0.15 | 0.63 |
| | Ball Vel X | Gaze X | -0.18 | 0.58 |
| | | Head X | 0.36 | 0.25 |
| | | Eye X | 0.17 | 0.61 |
| | | Gaze-ball X | -0.26 | 0.41 |
| | | Head-ball X | 0.14 | 0.66 |
| | Ball Vel Z | Gaze Y | -0.50 | 0.10 |
| | | Head Y | -0.26 | 0.41 |
| | | Eye Y | -0.46 | 0.14 |
| | | Gaze-ball Y | -0.46 | 0.13 |
| | | Head-ball Y | -0.13 | 0.70 |
| 30-40 | Ball Posi X | Gaze X | 0.20 | 0.54 |
| | | Head X | -0.04 | 0.89 |
| | | Eye X | -0.18 | 0.57 |
| | | Gaze-ball X | 0.01 | 0.96 |
| | | Head-ball X | 0.04 | 0.91 |
| | Ball Posi Z | Gaze Y | -0.13 | 0.68 |
| | | Head Y | 0.18 | 0.58 |
| | | Eye Y | -0.09 | 0.78 |
| | | Gaze-ball Y | -0.14 | 0.66 |
| | | Head-ball Y | -0.17 | 0.60 |
| | Ball Vel X | Gaze X | -0.23 | 0.48 |
| | | Head X | 0.49 | 0.10 |
| | | Eye X | 0.47 | 0.13 |
| | | Gaze-ball X | -0.50 | 0.09 |
| | | Head-ball X | 0.21 | 0.51 |
| | Ball Vel Z | Gaze Y | 0.48 | 0.12 |
| | | Head Y | -0.44 | 0.15 |
| | | Eye Y | -0.48 | 0.12 |
| | | Gaze-ball Y | -0.34 | 0.28 |
| | | Head-ball Y | 0.12 | 0.72 |
| 40-50 | Ball Posi X | Gaze X | 0.20 | 0.54 |
| | | Head X | -0.10 | 0.76 |
| | | Eye X | -0.15 | 0.63 |
| | | Gaze-ball X | -0.08 | 0.81 |
| | | Head-ball X | 0.02 | 0.95 |
| | Ball Posi Z | Gaze Y | -0.14 | 0.66 |
| | | Head Y | 0.09 | 0.78 |
| | | Eye Y | -0.32 | 0.31 |
| | | Gaze-ball Y | -0.29 | 0.39 |
| | | Head-ball Y | 0.18 | 0.58 |
| | Ball Vel X | Gaze X | -0.16 | 0.62 |
| | | Head X | 0.43 | 0.17 |
| | | Eye X | 0.47 | 0.12 |
| | | Gaze-ball X | -0.39 | 0.21 |
| | | Head-ball X | 0.33 | 0.29 |
| | Ball Vel Z | Gaze Y | -0.30 | 0.35 |
| | | Head Y | -0.06 | 0.85 |
| | | Eye Y | -0.48 | 0.12 |
| | | Gaze-ball Y | -0.48 | 0.11 |
| | | Head-ball Y | 0.21 | 0.52 |
| 50-60 | Ball Posi X | Gaze X | -0.01 | 0.97 |
| | | Head X | 0.03 | 0.92 |
| | | Eye X | -0.09 | 0.78 |
| | | Gaze-ball X | 0.01 | 0.98 |
| | | Head-ball X | 0.07 | 0.83 |
| | Ball Posi Z | Gaze Y | 0.35 | 0.27 |
| | | Head Y | 0.02 | 0.96 |
| | | Eye Y | 0.38 | 0.22 |
| | | Gaze-ball Y | 0.26 | 0.42 |
| | | Head-ball Y | 0.29 | 0.36 |
| | Ball Vel X | Gaze X | -0.40 | 0.20 |
| | | Head X | 0.55 | 0.07 |
| | | Eye X | 0.55 | 0.06 |
| | | Gaze-ball X | -0.37 | 0.23 |
| | | Head-ball X | 0.51 | 0.09 |
| | Ball Vel Z | Gaze Y | 0.28 | 0.37 |
| | | Head Y | -0.09 | 0.79 |
| | | Eye Y | -0.56 | 0.06 |
| | | Gaze-ball Y | 0.37 | 0.23 |
| | | Head-ball Y | 0.29 | 0.36 |
| 60-70 | Ball Posi X | Gaze X | 0.14 | 0.66 |
| | | Head X | -0.01 | 0.99 |
| | | Eye X | 0.15 | 0.63 |
| | | Gaze-ball X | -0.02 | 0.96 |
| | | Head-ball X | 0.15 | 0.64 |
| | Ball Posi Z | Gaze Y | 0.33 | 0.30 |
| | | Head Y | 0.09 | 0.78 |
| | | Eye Y | -0.49 | 0.11 |
| | | Gaze-ball Y | 0.04 | 0.90 |
| | | Head-ball Y | 0.34 | 0.27 |
| | Ball Vel X | Gaze X | -0.18 | 0.59 |
| | | Head X | 0.36 | 0.25 |
| | | Eye X | 0.51 | 0.09 |
| | | Gaze-ball X | -0.09 | 0.79 |
| | | Head-ball X | 0.34 | 0.28 |
| | Ball Vel Z | Gaze Y | -0.29 | 0.37 |
| | | Head Y | -0.20 | 0.54 |
| | | Eye Y | -0.54 | 0.07 |
| | | Gaze-ball Y | 0.25 | 0.44 |
| | | Head-ball Y | -0.41 | 0.18 |
| 70-80 | Ball Posi X | Gaze X | 0.14 | 0.67 |
| | | Head X | 0.09 | 0.79 |
| | | Eye X | 0.15 | 0.63 |
| | | Gaze-ball X | 0.10 | 0.76 |
| | | Head-ball X | 0.05 | 0.88 |
| | Ball Posi Z | Gaze Y | -0.13 | 0.69 |
| | | Head Y | -0.28 | 0.37 |
| | | Eye Y | -0.57 | 0.05 |
| | | Gaze-ball Y | 0.51 | 0.09 |
| | | Head-ball Y | -0.34 | 0.28 |
| | Ball Vel X | Gaze X | -0.38 | 0.22 |
| | | Head X | 0.38 | 0.22 |
| | | Eye X | 0.29 | 0.37 |
| | | Gaze-ball X | -0.11 | 0.74 |
| | | Head-ball X | 0.07 | 0.84 |
| | Ball Vel Z | Gaze Y | -0.23 | 0.47 |
| | | Head Y | -0.18 | 0.57 |
| | | Eye Y | -0.47 | 0.13 |
| | | Gaze-ball Y | -0.01 | 0.97 |
| | | Head-ball Y | -0.24 | 0.46 |
| 80-90 | Ball Posi X | Gaze X | 0.25 | 0.44 |
| | | Head X | 0.06 | 0.84 |
| | | Eye X | 0.22 | 0.50 |
| | | Gaze-ball X | 0.13 | 0.68 |
| | | Head-ball X | -0.08 | 0.80 |
| | Ball Posi Z | Gaze Y | -0.34 | 0.28 |
| | | Head Y | -0.35 | 0.26 |
| | | Eye Y | -0.43 | 0.17 |
| | | Gaze-ball Y | -0.10 | 0.76 |
| | | Head-ball Y | -0.18 | 0.59 |
| | Ball Vel X | Gaze X | -0.42 | 0.17 |
| | | Head X | 0.56 | 0.06 |
| | | Eye X | 0.48 | 0.12 |
| | | Gaze-ball X | 0.00 | 1.0 |
| | | Head-ball X | 0.25 | 0.43 |
| | Ball Vel Z | Gaze Y | -0.20 | 0.53 |
| | | Head Y | -0.21 | 0.51 |
| | | Eye Y | 0.54 | 0.07 |
| | | Gaze-ball Y | -0.27 | 0.40 |
| | | Head-ball Y | 0.11 | 0.74 |
| 90-100 | Ball Posi X | Gaze X | -0.15 | 0.63 |
| | | Head X | 0.23 | 0.48 |
| | | Eye X | 0.26 | 0.41 |
| | | Gaze-ball X | 0.26 | 0.41 |
| | | Head-ball X | -0.21 | 0.52 |
| | Ball Posi Z | Gaze Y | -0.45 | 0.15 |
| | | Head Y | -0.45 | 0.14 |
| | | Eye Y | -0.47 | 0.13 |
| | | Gaze-ball Y | -0.08 | 0.80 |
| | | Head-ball Y | -0.25 | 0.43 |
| | Ball Vel X | Gaze X | -0.51 | 0.09 |
| | | Head X | 0.49 | 0.11 |
| | | Eye X | 0.54 | 0.07 |
| | | Gaze-ball X | 0.31 | 0.32 |
| | | Head-ball X | -0.05 | 0.89 |
| | Ball Vel Z | Gaze Y | 0.13 | 0.69 |
| | | Head Y | 0.27 | 0.40 |
| | | Eye Y | 0.10 | 0.78 |
| | | Gaze-ball Y | -0.24 | 0.46 |
| | | Head-ball Y | 0.10 | 0.75 |
